# Free for all, or free-for-all? A content analysis of Australian university open access policies

**DOI:** 10.1101/2021.08.20.457045

**Authors:** Simon Wakeling, Danny Kingsley, Hamid R. Jamali, Mary Anne Kennan, Hamid Jamali, Maryam Sarrafzadeh

## Abstract

Recent research demonstrates that Australia lags in providing open access to research outputs. In Australia, while the two major research funding bodies require open access of outputs from projects they fund, these bodies only fund a small proportion of research conducted. The major source of research and experimental development funding in Australian higher education is general university, or institutional, funding, and such funds are not subject to national funder open access policies. Thus, institutional policies and other institutional supports for open access are important in understanding Australia’s OA position. The purpose of this paper is, therefore, to understand the characteristics of Australian institutional open access policies and to explore the extent they represent a coherent and unified approach to delivering and promoting open access in Australia. Open access policies were located using a systematic web search approach and then their contents were analysed. Only half of Australian universities were found to have an open access policy. There was a wide variation in language used, expressed intent of the policy and expectations of researchers. Few policies mention monitoring or compliance and only three mention consequences for non-compliance. While it is understandable that institutions develop their own policies, when language is used which does not reflect national and international understandings, when requirements are not clear and with consequences, policies are unlikely to contribute to understanding of open access, to uptake of the policy, or to ease of transferring understanding and practices between institutions. A more unified approach to open access is recommended.

## Introduction

In the context of scholarly communication, Open Access (OA) refers to the principle that research outputs should be freely and openly available for all users in a form which is “digital, online, free of charge, and free of most copyright and licensing restrictions” (Suber, 2012; p. 14). OA has been shown to have significant and tangible social and economic benefits (Tennant et al., 2016). Ensuring that access to published research outputs is equitable and cost effective is an important challenge for the higher education sector, and for Australian society at large (CAUL & AOASG, 2019).

The OA landscape is complex and multi-faceted. As Pinfield et al. have shown (2020), multiple dimensions operating at different levels involve many different actors combining in intricate ways. Advancing OA performance therefore requires the formulation and implementation of a range of strategies and processes carefully designed to address behavioural, technical and cultural issues. To take the example of just one geographic region – Europe – the last 10 years have seen a huge range of projects and policy initiatives designed to promote the uptake of OA publishing rates. The European Commission published their Guidelines on Open Access to Scientific Publications and Research Data in Horizon 2020 in 2016 (European Commission, 2016), and the associated Horizon 2020 programme represented a EUR30 billion investment in research and innovation between 2018-2020, under which “each beneficiary must ensure open access to all peer-reviewed scientific publications relating to its results” (European Commission, n.d.a). More recently, of course, Plan S has been launched (cOAlition S, 2021), a far reaching and somewhat controversial initiative designed to ensure that the outputs of publicly funded research be made available in Open Access form.

Individual European countries are also increasingly developing national approaches to open access and, more broadly, open research. Recent years have seen many countries propose national policies and strategies related to OA, for example Sweden (Swedish Research Council, 2015), Ireland (National Open Research Forum, 2019), and Finland (National Steering Group on Open Science and Research, 2020). In the UK the Research Councils UK open access policy (UK Research and Innovation, 2021), which took effect from 1 April 2013, and the Higher Education Funding Council for England (HEFCE) policy relating to articles submitted to the Research Excellence Framework (HEFCE, 2019), combined with the existing Wellcome Trust open access policy (Wellcome Trust, n.d.) has resulted in a significant shift to open access.

These national and supranational policies and frameworks are complemented by institutional polices. Recent research has shown that the European higher education sector has a sophisticated approach to institutional policy development and implementation, with 91% of institutions either having or developing an Open Science policy – a notable figure, given that such policies go beyond the boundaries of open access specifically, and embrace the broader concept of open science (Morais et al., 2021). The overall effect has been striking, with European countries in leading positions in rankings of global OA performance (European Commission, n.d.b).

In contrast, Australia, once a leader in global OA efforts, is now “lagging behind”, and there are urgent calls for open scholarship to be a national priority (CAUL & AOASG, 2019; Foley, 2021). Recent research evaluating international OA performance levels has shown that “universities in Oceania … lag behind comparators in Europe and Latin America” (see **Error! Reference source not found.**) (Huang et al., 2020). There is also increasing awareness that policies, and particularly OA mandates, are a crucial means of driving up OA adoption levels (Larivière & Sugimoto, 2018). In contrast to the European examples cited above, there is no overarching open access position in Australia. While major national funders such as the Australian Research Council (ARC) and the National Health and Medical Research Council (NHMRC) have policies requiring research to be made available OA, more than 30% of outputs from such funded projects fail to comply with this mandate (Kirkman & Haddow, 2020), and indeed these funders represent only 14.6% of all higher education R&D funds (Australian Bureau of Statistics, 2020). The major source of research and experimental development funding in Australian higher education organisations is general university, or institutional, funding (56%), and such funds are not subject to national funder OA mandates (Australian Bureau of Statistics, 2020).

The *Australian Code for the Responsible Conduct of Research* requires universities to develop and maintain “policies and mechanisms that guide and foster the responsible publication and dissemination of research” (Australian Government, 2018; p. 2). Since there is also a requirement that “institutions should support researchers to ensure their research outputs are openly accessible in an institutional or other online repository, or on a publisher’s website” (p. 3), it seems clear that institutional OA policies have an important role to play in driving OA performance. However, regional comparisons published by the Curtin Open Knowledge Initiative (COKI), which are dynamically updated over time (Figure 1), clearly indicate that institutions in Oceania, where Australia is located, perform relatively poorly in terms of open access (for the dynamic graphic see https://storage.googleapis.com/oaspa_talk_files/country_scatter.html).

**Figure 1:**
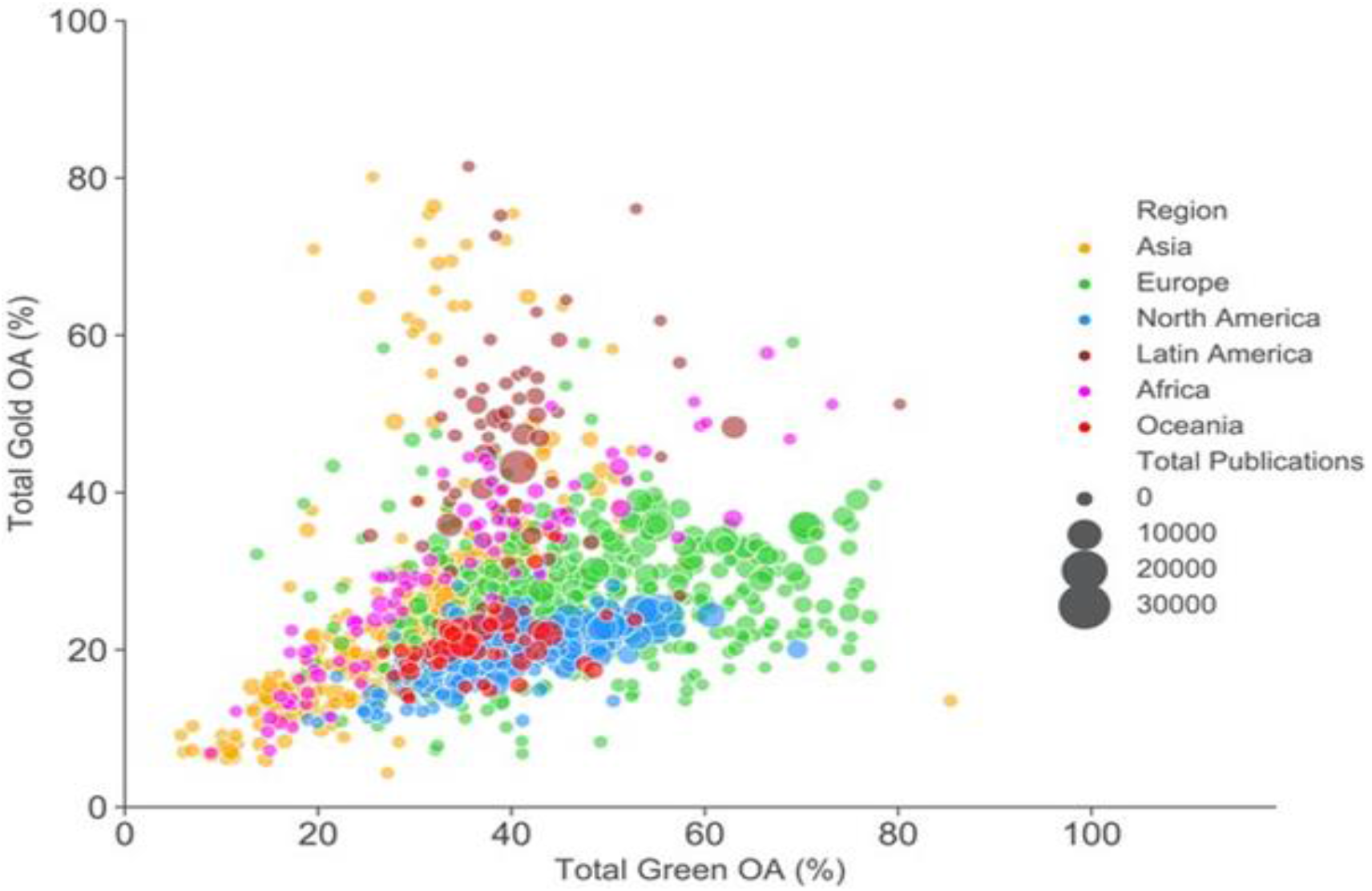
Comparing the level of gold and repository-mediated open access of individual universities - https://storage.googleapis.com/oaspa_talk_files/country_scatter.html

It is possible that one reason for this relatively poor performance is that the policy framework at an Australian institutional level does not effectively enough encourage or support OA. While in Australia the responsibility for managing open access policies is placed with institutions, to the best of the authors’ knowledge to date there has not been a detailed analysis of open access policies in Australian universities. This study aims to provide a content analysis of the open access policy landscape in Australia, considering key aspects of open access policies, including the means by which open access is achieved, the timing of the deposit of work into a repository and whether this differs to the timing of when it is required to be made openly available, the provision or otherwise of funds to support open access publishing (and whether there are restrictions around these funds), and other aspects of the policies. The stated intention and purpose of the policies is also a question of interest.

In particular this project sought to answer the following research questions:

- What are the characteristics of Australian institutional OA policies, in terms of content, intent and compliance mechanisms?
- To what extent do Australian university OA policies represent a coherent, unified approach to delivering and promoting OA in Australia?

### Literature review

There have been some prior efforts to identify OA policies at Australian Universities. Kingsley (2011) noted that in 2011 seven Australian Universities had OA “mandates”, but argued that several of these institutions were in fact “encouraging” rather than mandating OA. A later analysis by the same author found that as of January 2013, nine universities either had, or were in the process of implementing, OA mandates (Kingsley, 2013). Callan (2014) identified 12 institutions with OA policies in her review of the Australian OA landscape.

More broadly, a number of previous studies have reported individual institutions’ experiences of developing and implementing OA policies, both in Australia and internationally. Cochrane & Callan (2007) describe the development, implementation and impact of an eprint deposit mandate at Queensland University of Technology (QUT), describing collaboration between the University Research Office and Library, and the advocacy work required to obtain buy-in from research staff. Soper (2017) similarly outlines the background to the passing of an OA policy at Florida State University, with a rich description of institutional politics and advocacy efforts, and discussion of processes and systems developed to support its implementation. Kern & Wishnetsky (2014) highlight the role of the library in the development of an OA policy at Allegheny College, noting that an holistic effort was required encompassing advocacy and systems development, and emphasising the importance of institutional buy-in. Orzech & Meyers (2020) also outline the background to enacting an OA policy at several campuses of the State University of New York. They also highlight several key characteristics of institutional OA policies, including either mandatory or voluntary deposit, and opt-in or opt-out approaches. Otto & Mullen (2019) focus on the impact of the implementation of an OA policy at Rutgers, describing a significant increase in self-depositing practices while noting some confusion among researchers about which version of a manuscript to deposit. They also address systems, processes and strategies developed to encourage and facilitate self-archiving. Hoops & McLaughlin (2020) likewise address the systems side of OA policy implementation, describing the development process and functionality of an in-house application designed to support researcher depositing, while Kipphut-Smith (2014) presents the workflows developed to support the OA policy at Rice University. Saarti et al. (2020) report a survey of researchers undertaken to gauge attitudes and perceptions of open scholarly practices in the context of developing and implementing open science policies at the University of Eastern Finland. They note that a wider culture of open scholarship is only at an early stage, with associated challenges for instigating change through policy.

Beyond the institutional policy setting, earlier work has also focused on the nature and impact of national and funder policies. Crowfoot (2017) analyses the OA policies of the funding bodies who are members of the Science Europe association, noting some variation in the alignment of policies, particularly regarding embargoes, and positions on gold OA options. She also notes potential issues with monitoring and compliance mechanisms. Huang et al. (2020) report a detailed analysis of international OA performance at an institutional level, and link these performance levels to the various policy interventions at a national and funder level. In the UK, Awre et al. (2016) undertook a crowd-sourced review of OA policies as part of a broader JISC-funded project investigating OA support mechanisms. They report the development of column headings for a publicly available spreadsheet, with the purpose of crowd-sourcing a content analysis of OA policies affecting UK researchers. Finally, Kirkman & Haddow (2020) report a bibliometric study of Australian OA performance, with a particular focus on compliance with Australian funder OA policies, finding a compliance rate of just over two-thirds (67.3%).

Several recent publications relate to OA policy design, and highlight specific criteria that OA policies should include in order to maximise compliance and OA performance. Swan et al. (2015) conclude their analysis of the OA policy landscape with a series of recommendations. Specifically, they note that the OA deposit of research outputs should be mandatory, that deposit waivers should not be supported, that the depositing of research outputs should be linked to research evaluation, and that a requirement for “immediate” deposit is preferable. Morrison et al. (2020) produced a report for SPARC focusing on publisher copyright policies, which included recommendations for how institutions can “support their authors in maximising their research reach and impact by enabling OA” (p. 36). As well as suggesting that institutions provide support and guidance to researchers on copyright and licensing issues, the report also notes that research institutions should “work with publishers, funders and OA advocacy bodies to adopt standardised language when describing policy positions on copyright ownership and licensing” (p. 34) and “ensure standardised language is used by research offices, university libraries and academic schools when advising academic authors on OA copyright retention and reuse licence” (p. 36). Work by Larivière & Sugimoto (2018) echoes some of these issues in the context of funder policies, noting that allowing authors to deposit research after publication leads to lower deposit rates, and that the most effective policies are those with established and effective enforcement processes, and proper supporting infrastructure.

Most relevant to this study are the small number of publications that report surveys of institutional OA policies. While no such work appears to have been conducted in Australia, there are several international examples. Bosman et al. (2021) present a “multidimensional framework” incorporating different aspects of open access (for example what research outputs are covered by policies, how and when OA should be delivered, copyright issues), the different actors involved, and the levels of support encompassed by the policies. The authors review institutional, national and funder OA policies affecting Dutch researchers, and note whether these policies meet the various criteria outlined in the framework. The analysis provides important insights into the broader OA landscape, but seeks to illustrate the utility of the framework rather than undertake a granular content-based comparative analysis of the policies under review.

Fruin & Sutton (2016) report the results of a questionnaire relating to OA policies at US institutions. The questionnaire explored the rationale, processes and content of the institutions OA policies, and was directed only at institutions which have implemented, were in the process of implementing, or had attempted but failed to implement an OA policy. Of the 51 institutions which responded, 41% identified that their policy required deposit in an institutional repository (IR), while 14% characterised their policy as *merely a statement of encouragement to publish OA*, and 10% as asking faculty to *opt-in to the practice of self-archiving and open access*. Most policies were found to incorporate waivers for authors on request, and without any explanation being required. In the “vast majority” of cases, OA policies were found to originate from and be driven and managed by libraries. The questionnaire also sought to understand the arguments used to support the principles of OA, with the most commonly used justifications relating to author rights retention, access for all, and the goal of ensuring that publicly-funded research was publicly available. Perhaps the most significant findings relate to policy waivers and embargoes. The majority of institutions (70%) were found to respect publisher embargoes, with 10% not observing them, and 20% doing so only in certain cases. As the authors note, “the decision to honor these embargoes is typically an element of the implementation of the policy, which is usually led by the library as manager of the institutional repository” (p. 481).

Similarly, Duranceau & Kriegsman (2015) surveyed Coalition of Open Access Policy Institutions (COAPI) members about their OA policy development and implementation. From their analysis they identify four models of OA policy implementation: *systematic recruitment* (i.e. the library collecting publication metadata and using this to “request and acquire” publications from faculty), *targeted outreach* (i.e. the library targeting specific departments of faculties who are seen to be more “receptive” to OA), *faculty profiling* (i.e. use of a tool or system by which researchers can submit bibliographic metadata and upload outputs), and *harvesting* (i.e. the copying of research outputs from other repositories or publisher sites). The authors suggest that these approaches are potentially sequential, with institutions potentially moving towards a harvesting model as infrastructure and policy development allows.

Examining European OA policy alignment based on an analysis of ROARMAP (an international registry of self-reported OA policies), European funder policies and 365 institutional policies, Hunt & Picarra (2016) found that 63% of institutional policies mandated the deposit of research items, with 96% of policies specifying a repository as the locus of deposit. However, only 38% of institutional policies were found to include a requirement to make that deposit open access. The study also included analysis of deposit waivers, with 18% of institutional policies found to support waivers, and 21% found to not support them, with the remainder of the policies not specifying or not applicable. Examining the timeframe for making research outputs OA, 43% of institutional policies were found not to mention this, while 45% of policies specified either *when the publisher permits* or *by the end of policy-permitted embargo*. Only a very small number of policies were found to specify *acceptance date* (3%), *publication date* (3%) or *as soon as deposit is completed* (1%). It is important to note that a key finding of this study was that a significant number of institutional policies “do not specify or mention some essential elements which are critical to promoting a strong, effective policy” (p. 4), particularly relating to the time period for making outputs OA, length of embargo periods, ‘Gold’ OA publishing options, and a clear link to research evaluation (for example a note that noncompliance can impact the research assessment process).

From the early days of OA to more recent times the literature asserts how the various actions of government, funders and institutions (such as providing finance and supporting infrastructure, recognising and rewarding, legalising and promoting, setting as setting policy and stating as a goal), are all interrelated and reinforce each other in the uptake of OA. In this paper we analyse the content and intent of Australian institutional OA policies, as a means of better understanding how they might contribute (or otherwise) to the uptake of OA in Australia.

## Method

The study applied a content analysis approach. In order to identify institutions we used the Australian Government’s List of Australian Universities (https://www.studyinaustralia.gov.au/English/Australian-Education/Universities-Higher-Education/list-of-australian-universities) that includes 42 universities. We initially consulted the Registry of Open Access Repository Mandates and Policies (ROARMAP) to identify which of the 42 institutions have formal OA policies. However, it became apparent that whilst a useful resource, ROARMAP does not contain up to date information for some institutions. We therefore conducted our own systematic search of institutional websites in order to identify policies. This process first involved searching university websites to locate their policy libraries. Within the policy libraries, further searches were conducted using the following keywords:

- “Open access”
- “Publication”
- “Authorship”
- “Research”
- “Dissemination”
- “Theses”
- “Intellectual Property”

The policies, procedures or guidelines that were shown as a result of these searches were reviewed, and policies relevant to OA were downloaded and saved. Where institutions’ policy libraries did not appear to have any policies, procedures or guidelines that referred to Open Access, and for institutions without a searchable policy library, the university website was searched with the term ‘Open Access’ to identify any other relevant documentation.

The resulting documentation was reviewed, and the 42 institutions were then subjected to an initial classification of policy scope. A key question that informed this analysis related to how to define a “policy”. The language used in policy development is very specific. Terms such as policies, procedures, guidelines and others have particular meanings, and these are articulated in some instances (University of Queensland, 2020). National governance frameworks, such as those provided by the Tertiary Education Quality and Standards Agency, also emphasise the importance of formal policies. For the purposes of this study we therefore defined a formal OA policy as a document with the terms “open access” and “policy” in the title, **and** which was located either in the institution’s policy library, or elsewhere on the main university website. A second category consisted of institutions with policies that mention or relate to OA, but as part of a document with a broader scope (e.g. “Research publication policy”, “Research Repository Policy”). The third category consisted of institutions without a formal OA policy, but which provided less formal OA guidelines, principles or procedures document. These documents were sometimes found to be published on LibGuide sites. The final category related to institutions without any policy, formal or informal, relating to OA. Carnegie Mellon University was excluded from the study at this point, as the relevant policies were found to originate from the US parent institution, rather than be developed in Australia. This left a total of 41 universities.

This process resulted in identification of 20 (48.8%) universities which have a formal OA policy (**Error! Reference source not found.**). Eleven (26.8%) institutions were found not to have OA policies, but to have other policies that reference OA, while five (12.2%) universities without formal OA policies instead have other OA-specific documents titled principles, procedures or guidelines. Nine universities (22.0%) were found to have no policies, procedures or guidelines related to OA.

**Table 1:** The status of Open Access policy implementation in Australian universities

|  | No. of universities |
| --- | --- |
| Institutions with OA policies | 20 (48.8%) |
| Institutions without OA policies, but with other policies that reference OA | 11 (26.8%) |
| Institutions without OA policies but with OA Principles, Procedures, or Guidelines | 5 (12.2%) |
| Institutions that have no Policies, Procedures, or Guidelines referencing OA | 9 (22.0%) |

For the purposes of this research we focused our analysis on the 20 formal OA policies identified during the initial classification process. Including other types of policy that mention OA, or non-formal OA policy documents, would have made the analysis impractically complex, and would have required comparing documents with very different purposes and scope. All formal OA policies were downloaded between November 2020 and January 2021 and subjected to analysis using a mix of checklist and content analysis. We note here that some institutions may have updated existing policies, or introduced new ones, since the data were collected, and we acknowledge that the results are indicative of the OA landscape at the time the research was conducted. In selecting which aspects of the Australian policies to analyse, ROARMAP data categories were consulted along with some literature (Awre et al., 2016; Bosman et al., 2021). The categories used in the various analyses were different in each context, and used different language to describe similar concepts. To create a list of categories for analysis we borrowed from these previous studies and sites, then combined and/or added categories, labelling them in ways that seemed relevant to the Australian context and where possible using language reflected in the policies. This process involved all the research team and much discussion and refinement of categories. For example, for what we categorise as *output types included in the policy*, ROARMAP refers to as *content types specified*, Bosman et al. (2021) refer to as *What is made open access*, and Awre et al. (2016) as *A description of the type of research output which the policy covers*. Following this process each document was examined for information in the following categories:

- Date of first version of the policy
- Date of most recent version of the policy
- Date of next scheduled review of the policy
- Responsible office/Policy owner
- Definition of OA
- To whom the policy applies
- Output types included in the policy
- Timing for depositing outputs
- Role of the library
- The language used to describe responsibilities
- Exceptions to the policy
- Consequence for non-compliance
- Intent of the policy
- If and how funding for paying Article Processing Charges is covered in the policy

Some of the categories listed above were simple fact checking, but most items required more rigorous content analysis and categorisation. The categories (for instance categories for exceptions) were developed inductively by the researchers and were discussed and refined by the research team. The information collected for each policy was checked by at least two researchers to ensure its accuracy. All information and coding was recorded in a spreadsheet that is publicly available here: https://doi.org/10.6084/m9.figshare.15595572.v1.

The twenty OA policies were analysed in relation to their statements on paying for publication by creating a table that included any text copied from the policies that related to this issue. The text was then analysed for statements relating to hybrid publishing, any position on green or gold open access, whether funds were mentioned and if so whether any caveats were placed against those funds. We were aware that the absence of a statement doesn’t mean endorsement of the opposite position.

An analysis of the intent of the policies was more complex. Again each policy was scanned for language that referred to the purpose of the policy, but additionally any information that referenced a ‘position’ on how to approach open access. Once this text was extracted into a separate table, the text was analysed for any terms that were repeated or distinct. This identified different approaches from institutions such as recommending the use of an author’s addendum, or that authors retain the copyright of their work. A related analysis considered the specific language around the ‘benefit’ of the policy which fell into clear categories including the ‘benefit to society’, ‘increasing access to research’ and ‘maximising the research impact’.

### Findings and analysis

#### Number and date of policies

As noted above, 20 out of 41 Australian universities were found to have active OA policies. This represents an increase on the 11 institutions found to have policies in 2013 (Kingsley, 2013). In our study, where the date of the first version of these policies was reported (19), the oldest policy was from 2003, and the second oldest was created in 2008. The two years of 2013 and 2014 were clearly a key time for policy creation as 10 policies were implemented during this period. Three of the policies were created in 2020. The median age of the policies was seven years (created in 2014). However, we must acknowledge the possibility that some institutions may have had older policies that were subsequently superseded, and so were not included in our analysis. Analysis of the *Date of next scheduled review of the policy* data shows that of the 19 policies with a stated review date, ten show a historic date, with eight of these being pre-2020.

#### To whom the policy applies

Variation was found among OA policies in terms of the statement defining the people to whom the policies apply. In some cases definitions were detailed and granular: “This policy applies to all staff (including conjoint and adjunct staff) and students undertaking research at UNSW, either full-time or part-time and applies to scholarly research outputs” (UNSW). Others were much more general: “This Policy applies across the University” (ANU). Statements regarding to whom the policy applies were analysed, and categorized into four groups: staff, students, affiliates, and contractors. The findings have been summarized in **Error! Reference source not found.**, which shows that all OA policies were found to apply to staff, and the vast majority to students and affiliates.

**Table 2:** To whom the policy applies

| Categories | No. of OA policies |
| --- | --- |
| Staff | 20 (100%) |
| Students | 18 (90%) |
| Affiliates | 15 (75%) |
| Contractors | 1 (5%) |

#### Ownership of the OA policies

As might be expected, each OA policy was found to name a person or persons with responsibility for the policy. There was a surprising lack of consistency in the terminology used to describe these individuals and groups, reflective perhaps of different governance and organisational structures and associated nomenclature across Australian universities. Common titles were Accountable Officer; Administrator; Approval authority; Contact officer; Governing authority; Policy Custodian; Policy owner; Policy sponsor; Reference Authority and Responsible officer. Each policy was found to mention at least one of these terms. In some policies, a number of different roles were mentioned. For example, the Australian Catholic University policy names an Approval authority, Governing Authority, and Responsible Officer.

In 13 policies (65%) a position in the library was identified as responsible for the policy. Various positions have been allocated in charge of policies. In most cases the Library Director or University Librarian was named. Eight policies identified a non-Library contact as the person responsible for the policy, these most often being pro or deputy vice-chancellors. These results are broadly consistent with Fruin & Sutton’s findings relating to US OA policies (2016), which found that OA policies were library led in the “vast majority” of cases.

#### Role of the library in OA policies

Five OA policies (25%) did not mention the library. For the remaining 15, the role of the library was most often related to the institutional repository, delivering assistance or advice, and supporting copyright compliance (**Error! Reference source not found.**). It is important to note here that Fruin & Sutton’s study of US institutional OA policies found that while university libraries typically played a major role in OA policy development, implementation and support, these roles were often not articulated in the policies themselves. It is therefore possible that our results are not truly representative of the library involvement with OA activities.

**Table 3:** The role of libraries in OA policies

| Library Role | No. of mentions |
| --- | --- |
| Repository | 10 (50%) |
| Assistance/Help/Advice | 9 (45%) |
| Copyright Compliance | 8 (40%) |
| Reporting | 2 (10%) |

#### Mentions of funders in OA policies

References to funders were found in all 20 policies. Analysis of these references, however, reveals significant differences in the relationships between funder and institutional OA policies (**Error! Reference source not found.**).

**Table 4:**
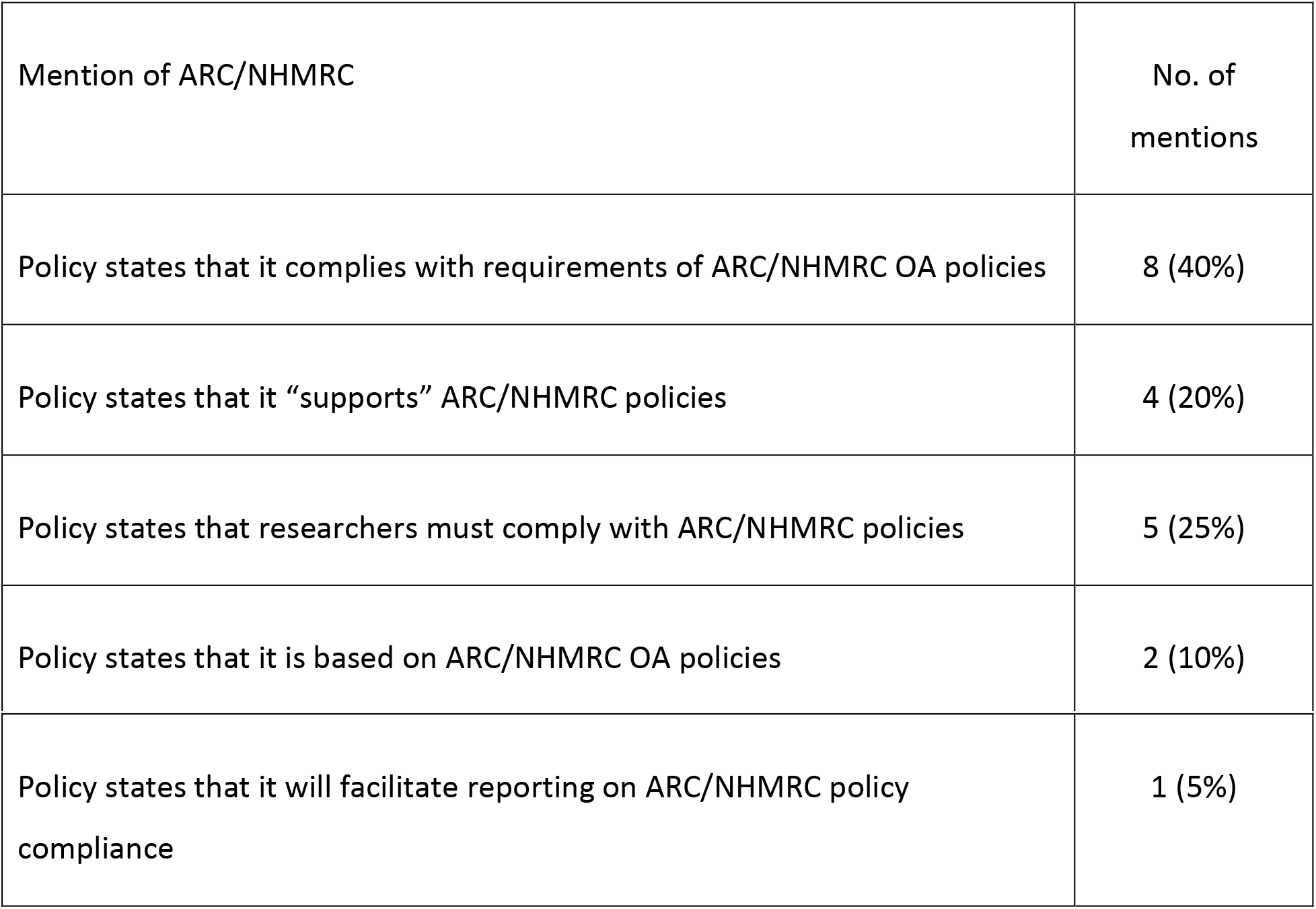
Mentions of funders in OA policies

| Mention of ARC/NHMRC | No. of mentions |
| --- | --- |
| Policy states that it complies with requirements of ARC/NHMRC OA policies | 8 (40%) |
| Policy states that it “supports” ARC/NHMRC policies | 4 (20%) |
| Policy states that researchers must comply with ARC/NHMRC policies | 5 (25%) |
| Policy states that it is based on ARC/NHMRC OA policies | 2 (10%) |
| Policy states that it will facilitate reporting on ARC/NHMRC policy compliance | 1 (5%) |

While one policy mentions funders only in the context of the policy being a tool to facilitate reporting and monitoring of compliance with funder policies, all the other policies incorporate funder OA policies is a substantial way. Funders were found to be most often mentioned in the context of institutions stating that their policies comply with funder requirements. The implication here is that institutional policies have been designed to ensure alignment with minimum funder requirements, such that researchers acting in accordance with the institutional policy will automatically comply with ARC and NHRMC policies. In addition, two policies – those of QUT and the University of Queensland – while not specifically mentioning compliance, explicitly state that they are based on the requirements of ARC/NHMRC policies. In contrast, five policies include specific requirements for authors to comply with national funder OA policies, in addition to the requirements outlined elsewhere in the policies, implying that adhering to the standard policy requirements alone would not ensure compliance with funder policies. In the remaining four cases, policies are less clear about the relationship between institutional and funder policies, stating only that they “support” funder policies, and including no specific requirements to adhere to funder policies.

As noted above, the major Australian national research funders (ARC and NHMRC) represent only 14.6% of all Higher Education Research & Development funds (Australian Bureau of Statistics, 2020). Nonetheless, their OA policies have clearly had a strong influence on institutional OA policy development. This in turn suggests that stronger funder policies will result in stronger institutional policies, and improved OA performance nationally. This is supported by evidence from recent literature which has highlighted the effects of funder policies on national OA performance levels in countries such as the US, UK, Canada and the Netherlands (Larivière & Sugimoto, 2018; Huang et al., 2020; Robinson-Garcia et al., 2020)

#### Definitions of Open Access

As might be expected in a formal policy document, a very high proportion of university OA policies (90%) were found to include a definition of OA. It was expected that policy definitions would reflect commonly understood definitions of OA. While the majority of policies had a definition of OA, very few of the definitions were the same. Definitions were shared only in two cases, each by two universities. It would be interesting to know if there was collaboration on policy development in these cases.

Shared understandings are more likely to make it easier to implement OA within and between universities, at national and international levels, and to build community acceptance and understanding of OA models. Faced with the challenge of drafting a definition of OA for inclusion in an OA policy, we might expect authors to borrow from definitions found in well-known OA initiatives or related policies, such as the Budapest Open Access Initiative (BOAI) (2002), Berlin Declaration on Open Access to Knowledge in the Sciences and Humanities (Berlin) (2003), or UNESCO (n.d.). Given the Australian context, we might also expect to see text based on the AOASG (2019 and based on the Budapest and Berlin declarations) or even the ARC (2017) and NHMRC (2018) definitions. Searches in both Google and Google Scholar revealed that most definitions covered aspects from the above definitions, but only two referenced the sources of their definitions, one of these referencing AOASG (the Australian Catholic University) and one BOAI (University of New South Wales).

The ARC/NHMRC definition covers reuse, licensing and attribution which are key concepts in understanding open access; however, most definitions used simplified language and some focused only on access, for example:

“Open Access means immediate, permanent, unrestricted, free online access to the fulltext of refereed research publications.” (University of New England and James Cook University)
“Open access: Allowing research outputs to be freely accessible to the general public;”

One made the definition local to their own organisation:

“Open access means free and unrestricted (electronic) access to [Institution] conducted research, articles and other scholarly outputs.” (La Trobe University)

Another made the definition only relate to green open access in a repository and did not reflect other OA options:

“Open Access means permanent, free online access to research and scholarly publications through a central repository on the public internet” (Southern Cross University)

These simplified definitions did not refer, for example, to reuse, licensing and attribution which are key concepts in understanding open access, misunderstandings of which may affect the likelihood of researchers to make their work open access (Zhu, 2017). While it is understandable that definitions are simplified for accessibility of understanding, definitions which do not reflect national and international understandings or miss important aspects of open access are unlikely to contribute to understanding of OA, to uptake of the policy, or to ease of transferring understanding and practices between institutions. It is also important to note Morrison et al.’s (2020) recommendation that successful OA policies should use standardized and consistent language both within and across universities.

#### Exceptions

All 20 OA policies were found to specify various exceptions to the standard requirement that work be made available OA (**Error! Reference source not found.**). It should be noted here that the category creation was driven by the language used in the policy. It could be argued, for example, that *Publisher agreement* and *Copyright or licensing restrictions* are very closely related, but it was thought important to note the difference in language used in the policies. In practice this distinction typically referred to the difference between publisher embargo periods and other restrictions. For example:

“Where deposit of the full-text material, or dataset, is not possible due to publisher embargo, or is not permissible due to copyright or licensing restrictions” (Bond University)

Other exceptions included concerns related to commercial or cultural sensitivity, and confidentiality, although less than half of all OA policies specified these exceptions.

**Table 5:** Exceptions to the standard requirements in OA policies

| **Exception type** | **No. of Policies** |
| --- | --- |
| Publisher agreement<br>(including embargo) | 18 (90%) |
| Copyright or licensing<br>restrictions | 8 (40%) |
| Commercial sensitivity | 6 (30%) |
| Confidentiality | 6 (30%) |
| Cultural sensitivity | 3 (15%) |
| Legal obligations | 2 (10%) |
| Already OA material | 1 (5%) |
| Publisher version of non-<br>open access publications (i.e.<br>version of record) | 1 (5%) |
| Royalty payment or revenue | 1 (5%) |

It is instructive to consider these findings in light of the findings of previous studies. Fruin & Sutton’s study of US OA policies (2016) found that just 10% of institutional policies specified not observing publisher embargoes, with most policies incorporating waivers for authors, both results broadly consistent with our findings. Similar results were also found in Hunt & Picarra’s study (2016), with 21% of institutional policies not supporting any waivers to OA deposit. It seems clear that there is a general trend for institutional OA policies to explicitly respect publishers’ positions on OA, thus not following Swan et al.’s (2015) recommendation that such policies should not allow waivers.

#### Language used to describe the policy directive

Our content analysis also included identification of the language used in association with the OA directive. **Error! Reference source not found.** presents the most commonly found words used in association with the instructions to researchers, along with examples of the terms in context. In many cases the language is strong (“must”, “required”), implying a mandate even if this particular word is not included. We also note a distinction between whether policies use an active (“researchers will …”) or passive (research outputs must be made…”) form. Once again, the language used to describe the policy is varied, in contrast to Morrison’s recommendation (2020) that policy language should be standardised across institutions. One might also question the extent to which the language used constitutes a “mandate”, as recommended by Swan et al. (2015). Notably only one policy was found to use the word “mandate”.

**Table 6:** Language and action verbs used in association with OA directives

| Key word | No. of OA Policies | Example |
| --- | --- | --- |
| must | 5 (25%) | "... fulltext research outputs **must** be made openly available where ..." |
| will | 4 (20%) | "Researchers **will** ... make publications and data arising out of research openly available for re-use and citation." |
| requires/required | 3 (15%) | "The University **requires** all staff and students to deposit Research Outputs ... for the purpose of providing Open Access" |
| responsible/responsibility | 3 (15%) | "the following **responsibilities** are in place ... Secure where possible the immediate unrestricted access to publication" |
| is to be | 2 (10%) | "material ... **is to be** deposited in the University's open access institutional repository" |
| mandates | 1 (5%) | "The University **mandates** an open access approach ..." |
| should be | 1 (5%) | "Research Outputs **should be** forwarded to the University Library for deposit into the institutional repository ... if an open access version is not already available" |
| encourages | 1 (5%) | "The University **encourages** staff and students to ..." |

#### Compliance/consequences of breach of policy

Taken in isolation, the language associated with OA directives, as described above, might be considered relatively strong. It is important, however, to consider this language not only in terms of the associated exceptions to the policy, but also to the compliance and monitoring mechanisms associated with the policy, and the stated consequences of a failure to comply. We found that OA policies typically do not explain the monitoring processes. This is perhaps not surprising, as a detailed outline of such activity might reasonably be considered to be beyond the scope of a formal policy document. It is also important to recognise that the requirement to comply with University policies, and consequences of non-compliance, may be outlined in other formal documents (such as employment contracts and codes of conduct). It is notable, however, that only three of the twenty OA policies (15%) specifically state the consequences of a failure to comply. We note that in all three cases, the impact of the statement is softened by the inclusion of the word “may”:

“The University may commence applicable disciplinary procedures if a person to whom this policy applies breaches this policy (or any of its related procedures)” (Macquarie University)
“Non-compliance with this Policy may constitute research misconduct and/or general misconduct, which will be addressed in accordance with the University’s Enterprise Agreement and relevant disciplinary procedures.” (University of Adelaide)
“Breaches of this Policy may result in action being taken in accordance with the University Code of Conduct for Research.” (UNE)

That so few policies were found to explicitly discuss compliance monitoring is consistent with the findings of Hunt & Picarra (2016), who argued that “a significant number” of OA policies omit elements, including monitoring, and links to research assessment activities, that are “critical to promoting strong, effective policy”. Crowfoot (2017) has noted the issues with compliance and monitoring in the context of funder policies, while Larivière & Sugimoto (2018) have argued that the best policies are those which are effectively enforced. It is likely that compliance monitoring activities (of varying types) are undertaken at many institutions, without being specified in policies. Nonetheless we argue that specifying compliance activities, and consequences for breaches of policy, would strengthen OA policies.

#### Payment for publication and hybrid journals

We examined the specific statements used within policies that relate to paying for OA publication and these are reported in Figure 2. Detail of the positions taken on the payment of article processing charges (APCs) in subscription journals in particular (known as hybrid OA) are relevant, as charging authors an APC and readers a subscription for the same article has led to accusations of double dipping by these publishers (Phillips, 2020). A 2016 analysis of requirements of UK funders, and of US and UK library-run funds found there was a wide variety in the way the expectations around hybrid was expressed (Kingsley & Boyes, 2016). Given the Australian investment has historically focused on green open access (putting a copy of the work into repository), the assumption could be made that hybrid would be at odds with this strategy in Australia.

**Figure 2:**
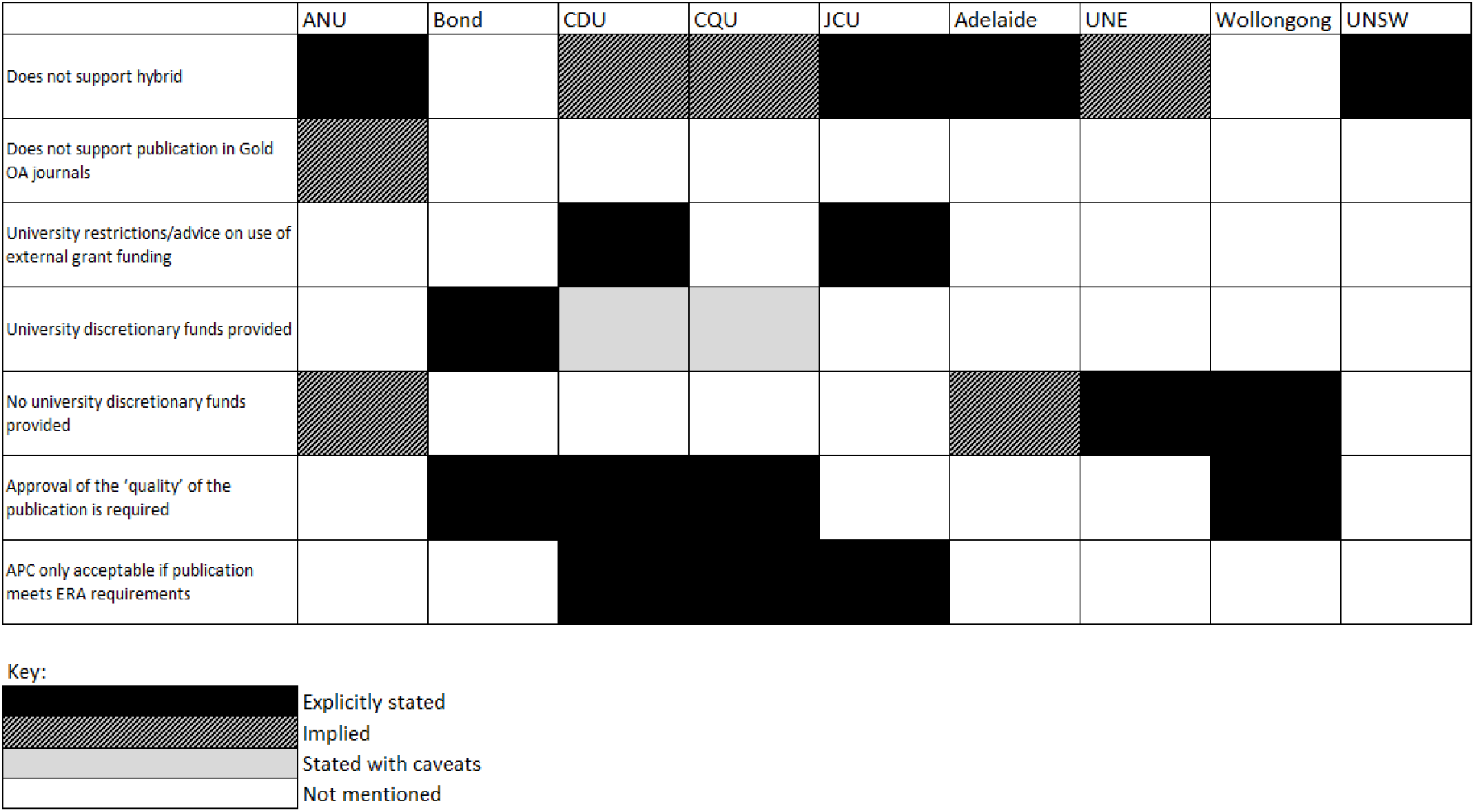
Analysis of the nine Australian university OA policies which mention paying for publication

In fact, the variation found in the UK and US policies was reflected in our analysis of Australian OA policies. Some of the policies are indeed clear on the question of hybrid. James Cook University says “Hybrid Open Access publication is not supported by this policy.” The University of Adelaide states: “The University recommends that researchers should avoid paying APCs to publish in Hybrid Journals”, and UNSW says: “UNSW discourages authors from paying Article Processing Charge (APC) fees to make outputs open access in an otherwise subscription-based (hybrid) outlet (sometimes called publisher double-dipping).” The Australian National University states it does not support hybrid, but the use of the word ‘or’ in the policy means it could be interpreted to say it does not support the payment of any article processing charges: “The University does not support the payment of article processing fees (APCs) or ‘hybrid’ fees (where an individual article is made available through payment of an article processing fee)”.

Neither Bond University nor the University of Wollongong mention hybrid at all, but have opposing positions on paying for publication. Bond makes no restrictions or suggestions on paying for publication: “Use of external grant funding or discretionary University, Faculty, or Research Centre funding may be provided to cover the publishing costs i.e. Article Processing Charges, of accepted open access publications”. Wollongong on the other hand states: “The University maintains a position to not pay for the publishing of online research where possible.” However, the policy goes on to say: “Gold Open Access may be supported and funded at the faculty level where strategically or otherwise appropriate.” It does state: “The University supports a Green approach to Open Access” which could possibly be interpreted as being anti-hybrid, but it is not specific.

There are implied sanctions on hybrid in several policies that explicitly support ‘Gold Open Access Journals’ and which do not specifically mention hybrid. For example, Charles Darwin University uses the expression: “In a journal that is deemed to be Gold Open Access”. Central Queensland University says: “the publishing outlet is considered to be Gold Open Access” and the University of New England suggests researchers should seek funding: “if wishing to publish in an open-access journal”.

Perhaps more than any other element of our analysis, the findings relating to positions on paying for publication illustrate the lack of a consensus vision for OA in Australia. Given Morrison’s persuasive arguments for consistency and standardisation across institutions (2020), and the complexities associated with different visions for OA, this represents a significant issue for the delivery of OA in Australia.

#### Timing of deposit

There is evidence to demonstrate that the timing of a policy can make a difference to the compliance rate (Herrmannova et al., 2019). Only 13 of the 20 Open Access policies specify a timeframe within the policy, and there were found to be significant variations within these policies. The University of Melbourne and the University of Wollongong were found to specify the period of time within which the work should be deposited into the repository (within three months of publication and at the time of publication respectively). Australian National University has different requirements for deposit to the repository depending on the item in question, but for journal and conference publications, technical reports and other original, substantial works, the timing is: “within 3 months or as promptly as possible after publication”. Four universities specify deposit of work into the repository “upon” or “after” acceptance for publication: Edith Cowan University, Macquarie University, Queensland University of Technology and Australian Catholic University. These are interesting because managing embargoes can be challenging if a work is deposited prior to publication. Publisher embargoes relate to a period of time after publication, so institutions requiring deposit prior to publication must have systems in place to revisit these works at the time of publication to set the embargo. As Larivière and Sugimoto (2018) have noted, it is essential that OA policies are supported by proper technical infrastructure and process design, for example, to switch embargoed deposits to open once embargo periods are over.

This distinction between when something is deposited and when it is made openly available means in some instances a deposited item might not be made openly available for up to 24 months after deposit. For this reason, it is significant if a policy does stipulate when a work should be openly available, as distinct from simply when a work should be deposited. One example is the University of Adelaide which states: “Researchers are encouraged to avoid embargoes of greater than 12 months from date of publication. Where agreements do not allow outputs to be made Open Access within 12 months researchers should make reasonable attempts to negotiate this provision with the publisher.” University of South Australia aspires for deposit and open access with the text: “as soon as is practicable and not later than twelve months after publication”. The University of Sydney states “no later than 12 months after the date of publication”. Both University of New South Wales and the University of Queensland identify making work openly accessible “within twelve months of publication” but both have the caveat “or as soon as possible”. Western Sydney University is even more passive on pushing back on embargos, simply stating “as soon as possible”.

This inconsistency and lack of clarity about what is actually required in terms of deposit and openly accessible timeframes is a significant issue in relation to having a unified position and policy across the country. The variations in Australian institutional policies are consistent with findings from other studies, including Hunt and Picarra (2016), who found that 43% of European institutional policies failed to mention the timing of deposit, with most of the remainder using vague terms such as “when the publisher permits”.

#### Intent of policies

Considering the text used within a policy that refers to the intent or rationale behind the policy, a series of perspectives arise. By considering the different policy statements it is possible to identify if a policy supports particular positions on the question of open access. As is identified in **Error! Reference source not found.**, different policies make a statement about whether the university supports green or gold OA, and whether they have a position on hybrid journals. Some articulate a requirement about the retention of intellectual property and some offer options for managing this through an addendum. Some policies define the support the university will be providing and others are clear about the accountability of the effectiveness of the policy.

**Table 7:** Rationale used in policies in support of OA as a principle

|  | No. of OA Policies |
| --- | --- |
| Open benefits society | 8 (40%) |
| Increases access to the university's research | 7 (35%) |
| Maximises research impact | 6 (30%) |
| Infrastructure | 2 (10%) |
| Reproducibility | 2 (10%) |

When considering the motivation behind the policies, we identify three main categories: to increase the profile of the research of the university, to ensure the university research has a wide audience, or because open research benefits society. As with other aspects of our analysis, there is a clear lack of consistency in terms of the rationale used, with no single argument being adopted by more than half of the studied universities. Nine universities have a statement of intent within the policy which refers to a wider benefit beyond the institution. However, these vary considerably. The Australian Catholic University policy stands out because it is widely encompassing and quite specific in its stated purpose, referencing the Australian Deputy Vice Chancellors (Research) Committee mandated Findable, Accessible, Interoperable and Reusable (FAIR) <u>Statement</u>; intellectual, social, economic, cultural and environmental benefits; excellence, impact and engagement in research practice, including reproducibility and approaches that encourage collaboration and the transfer of knowledge between researchers and users of research in industry, government and the general community.

The remaining policies that are classified within the ‘open research benefits society’ vary in scope and aim. Some policies specifically refer to benefits to society, for example the University of Queensland policy aims to “ensure the results of research are made available to the public, industry and researchers worldwide, for the benefit of society” and the University of New South Wales states that: “Open Access publication enables us to share our capability in research and education effectively and equitably with global partners and stakeholders … Open Access supports the generation of new knowledge applied to solve complex problems, deliver social benefits and drive economic prosperity, locally, nationally and globally”. Others are more tangential in their reference to benefits, such as James Cook University which notes the policy is “recognising that knowledge has the power to change lives”.

Others are more focused on exchange of information with the public. For example, the University of Sydney states the policy “supports the University’s core values of engaged inquiry and mutual accountability by encouraging transparency, accountability, openness and the sharing of scholarly works and other research outputs with the research community and the public”. Both Macquarie University and the Australian National University refer to the open exchange of information as a “bedrock academic value”. Despite the general philosophy that publicly funded research should be publicly available, only Edith Cowan University and the University of New England specifically refer to “publicly-funded research”.

There are some significant differences between our findings, and those that emerged from Fruin & Sutton’s (2016) survey of US librarians. They found that author rights retention and the ensuring public access to publicly funded research were among the most commonly used arguments to support the principles of OA. Neither of these arguments featured in our analysis of Australian OA policies. It is worth noting, however, that Fruin & Sutton’s question related to the arguments used “in conversations with faculty and other constituents about why open access is important”, rather than the arguments used in the policies themselves. The different arguments used in these different contexts are potentially revealing, and perhaps merits further research.

## Discussion

### Prevalence and strength of Australian institutional OA policies

The Australian Code for the Responsible Conduct of Research devolves responsibility to institutions to develop and maintain OA policies (Australian Government, 2018), but the results this study demonstrates that only 20 (50%) of universities have a formal OA policy. This finding is particularly concerning given the wealth of evidence from around the world that confirms the positive impact of strong and consistent OA policies (Herrmannova et al., 2019; Huang et al., 2020; Larivière & Sugimoto, 2018; Rieck, 2019; Robinson-Garcia et al., 2020).

Funder mandates, institutional policies, grass-roots advocacy, and changing attitudes in the research community have contributed to the considerable growth in open access publishing during the last two decades (Huang, et al., 2020). While having an open access policy in place is important, the literature demonstrates that a policy alone is not enough to ensure the OA of research outputs. Clear language and processes for the enforcement of OA in policies is required to stimulate significant growth of OA. Our analysis found that none of the 20 existing Australian university OA policies mentioned monitoring of compliance, and only three specified any consequences for a failure to comply. While there may well be monitoring activities taking place at many Australian universities, it is clear that these are not widely publicised, and certainly not codified in policy documents. However, it has been shown that there is a clear link between compliance rates and clearly stated consequences for non-compliance (Larivière & Sugimoto, 2018). To give one example, the National Institutes of Health introduced an open access policy in 2005, requesting that funded authors deposit a copy of their publication in PubMed Central (Suber, 2008). In November 2005 it was reported that fewer than 3% of publications were being deposited and recommended that the policy become a requirement (NIH Public Access Working Group, 2005). A new policy was written, mandating deposit, applicable from April 2008 (National Institutes of Health, 2008). By the end of that year compliance was almost 50% (Poynder, 2009). In 2013, the NIH strengthened the policy further, in that it would: “delay processing of non-competing continuation grant awards if publications arising from that award are not in compliance with the NIH public access policy” (National Institutes of Health, 2012). This resulted in a ‘surge’ of deposits of papers, both current and retrospective (Van Noorden, 2013). Similarly, the Wellcome Trust has also strengthened their open access policy over time. Their policy was introduced in 2005 but compliance was only at 15% in 2007 (Wellcome Trust, 2014). In June 2012, when compliance was at 55%, the policy changed, requiring grant recipients to demonstrate compliance otherwise “the final payment on the grant will be withheld” and new grants would not be awarded (Wellcome Trust, 2012). Following the stronger requirement, by 2019 compliance was up to 95% (Wellcome Trust, 2019). This evidence should provide strong incentive to policy makers at all levels, including Australian Universities, to ensure that OA policies include meaningful consequences for compliance failures.

### Standardisation, consistency, and aligned intent

Our analysis clearly demonstrates enormous variation across OA policies. We found significant differences in the intent of policies; the definitions of OA; the arguments used to support it; requirements for the timing of OA deposits; positions on paying for publication; the language used to describe researcher responsibilities; the exceptions to OA requirements, and the role of libraries in both policy development and compliance and monitoring. While we recognise that institutions are independent entities with their own goals and priorities, and therefore some variation is perhaps inevitable, the overall picture is one of confusion and inconsistency. This is especially troubling given that Australian researchers do not work in institutional vacuums. Cross-institutional collaboration is commonplace in Australia (Luo et al., 2018), as is researcher mobility, with most academics working at multiple universities over the course of their careers (Bexley et al., 2011).

Our analysis demonstrating the wide variety of positions on, and language about, OA in Australian institutional policies is likely to suggest to researchers that OA is a fractured concept. This may be a result of OA policies being developed and owned by a range of roles and institutional areas which may have different priorities and account for some of the lack of standardisation in Australian university OA policies. While evidence has repeatedly shown that clear, consistent national or supranational positions on OA are the most effective way of maximising OA performance, our study suggests that Australian universities are a long way from achieving this. In the absence of clear national leadership it is vital that HE institutions themselves work collaboratively to build consensus around effective OA policy development. Further research in this area, in order to better understand the key stakeholders and organisational processes at play, and identify the appropriate mechanisms for collaboration, would undoubtedly be beneficial.

The stated intention of the policies analysed in this study differ markedly, ranging from increasing the impact of the institution’s research outputs, to increasing the profile of the institution, to improving society. None of these rationales are problematic in themselves, however the disparity across the policies indicates a lack of shared purpose. The policies serve different purposes within institutions. It is interesting that of all of the policies, only two mention the position that ‘publicly funded research should be publicly available’, which has been a longstanding justification for open access. For Australia to have a position on open access, reflecting activity in Europe, it will be necessary to come to an agreed position on why open access is needed.

### Clarity on APCs

Our analysis of the varied positions on paying for publication, whether as gold or hybrid, indicates that Australia is very far from putting forward a unified position on APCs. As well as the general and overarching point that this lack of consistency is confusing for researchers, this is particularly significant given the recent shift to what are becoming known as ‘transformative’ deals between academic libraries and publishers. These incorporate the costs of publication as well as the costs of subscriptions with a general aim of reducing overall costs or at least remaining cost neutral (Hinchcliff, 2019). In order to assess the value of a proposed deal to an institution, there is a pressing need to understand the institutional expenditure on APCs. In Australia, where APCs are mostly paid by individual grant holders and not centrally managed, it is difficult to determine the level of expenditure on them. Attempts to identify this figure date as far back as 2014 (Kingsley, 2014).

In Australia, group negotiations on content procurement are managed by the Council of Australian University Librarians (CAUL) through a CAUL consortium. In October 2019, CAUL announced the first transformative agreement for Australia and New Zealand with the UK-based Microbiology Society, which provides the university libraries the ability to pay a single “publish and read” fee for uncapped open access publishing in all of the Society’s journals by corresponding authors (CAUL, 2019).

During 2019 CAUL also commenced a project to design and implement a consistent process for collection and reporting of APCs (Cramond et al., 2019). As transformative deals become more common in the Australian landscape, the need to have clearer policies relating to APCs and a better understanding of APC expenditure at an institutional and national level becomes more urgent. This in turn requires institutions to develop consistent policies on funding APCs, and clearer guidance for researchers on their use.

### Timing of deposits

Only 13 of the 20 institutional OA policies were found to specify a deadline for deposit of papers into a repository. In many of those 13 we found inconsistency, and a conflation between instructions to ‘deposit work’ and ‘make the work openly accessible’ in the language of the policies. ‘Deposit’ and ‘make open’ are different actions and clear differentiation of the two in any OA policy would assist researchers. For example, if a policy states that a work must be deposited on publication, yet there is a publisher embargo on open OA, then the work must initially not be OA and only be available as a metadata-only record in the repository. The full work may only be made openly accessible once the embargo period is complete. It is essential that those responsible or drafting policy understand the challenges for researchers associated with understanding this complex space, and draft policy accordingly. As there is a positive correlation between requiring deposit closer to acceptance (rather than on or after publication) (Larivière & Sugimoto, 2018),compliance policies should ideally both stipulate when a work needs to be deposited into a repository, and when the work needs to be made OA. Thus there are significant operational implications of not clarifying this differentiation, and not noting when a deposited item is subject to a publisher embargo.

## Conclusion

In this article we have reported in detail the results of a content analysis of formal institutional open access policies at Australian universities. Just 20 of 40 Australian universities were found to have such a policy, despite the *Australian Code for the Responsible Conduct of Research* requiring universities to publish “policies and mechanisms that guide and foster the responsible publication and dissemination of research” (Australian Government, 2018). Within the 20 analysed policies we found extensive variation across a number of crucial areas, including paying for publication, deposit timing, and the intent and rationale underpinning the policies. In addition, we found only three policies which explicitly stated the consequences for noncompliance.

There is growing impetus towards the development of a national OA strategy in Australia. The new Australian Chief Scientist, Dr Cathy Foley, has indicated her support for a unified approach, a move welcomed by advocacy groups CAUL & AOASG, 2021). Our findings show just how vital consensus building and standardisation will be as part of this process. We suggest that there is an urgent need for collaborative, inclusive and detailed discussions involving a range of stakeholders, to ensure that there is not only a common goal, but a clearly defined framework - including consistent and effective institutional policy development and implementation - to achieve that goal.

